# EVscope: A Comprehensive Bioinformatics Pipeline for Accurate and Robust Analysis of Total RNA Sequencing from Extracellular Vesicles

**DOI:** 10.1101/2025.06.24.660984

**Authors:** Yiyong Zhao, Himanshu Chintalapudi, Ziqian Xu, Weiqiang Liu, Yuxuan Hu, Ewa Grassin, Minsun Song, SoonGweon Hong, Luke P. Lee, Xianjun Dong

**Author notes:** Corresponding author: Xianjun Dong.

## Abstract

**Motivation:** Extracellular vesicle (EV) RNA sequencing has emerged as a powerful approach for studying RNA biomarkers and intercellular communication. Nevertheless, the extremely low abundance, fragmented nature and ubiquitous tissue origin of EV RNAs, alongside potential contamination from co-isolated materials, such as free DNA and bacterial RNA, pose substantial analytical challenges. These complexities highlight a pressing need for a standardized, computational workflow that ensures robust quality control and EV RNA characterization.

**Results:** Here, we present EVscope, an open-source bioinformatics pipeline designed specifically for processing EV RNA-seq datasets. EVscope employs an optimized genome-wide expectation-maximization (EM) algorithm that significantly improves multi-mapping read assignment at single-base resolution by effectively leveraging alignment scores (AS) and local read coverage, specifically tailored for fragmented and low-abundance EV RNAs. Notably, EVscope uniquely generates EM-based BigWig files for downstream analysis, a capability currently unavailable in existing EM-based BigWig quantification tools. The pipeline systematically integrates 27 major steps, including quality control, analysis of library structure, contamination assessment, read alignment, read strandedness detection, UMI-based deduplication, RNA quantification, genomic DNA (gDNA) contamination correction, cellular and tissue source inference and visualization with a comprehensive HTML report. EVscope incorporates a comprehensive, updated annotation covering 19 distinct RNA biotypes, encompassing protein-coding genes, lncRNAs, miRNAs, piRNAs, retrotransposons (LINEs, SINEs, ERVs), and additional non-coding RNAs (tRNAs, rRNAs, snoRNAs). Furthermore, it leverages two highly balanced circRNA detection algorithms for robust circular RNA identification. Notably, a downstream module enables the inference of the tissue/cellular origins of EV RNAs using bulk and single-cell RNA-seq reference datasets. EVscope is implemented as a convenient, single-command Bash pipeline leveraging Conda-managed standard software packages and custom scripts, ensuring reproducibility and straightforward deployment.

**Availability and implementation:** Code, documentation, and tutorials are available at GitHub (https://github.com/TheDongLab/EVscope) and archived on Zenodo (https://zenodo.org/records/15577789).

## 1. Introduction

Extracellular vesicles (EVs) mediate intercellular communication primarily through the transfer of functional RNA molecules. These nanosized, membrane-bound structures are secreted by nearly all cell types and regulate numerous physiological and pathological processes (Yáñez-Mó, et al., 2015). Consequently, RNAs derived from EVs have emerged as promising biomarkers for various diseases (Kumar, et al., 2024), including cancer and neurodegenerative disorders, highlighting the growing importance of EV-RNA sequencing (RNA-seq) (Cheng and Hill, 2022). Nevertheless, technical challenges remain due to the inherent characteristics of EV RNAs, such as their extremely low abundance, high heterogeneity (Willms, et al., 2018), fragmented RNA, and susceptibility to contamination by genomic DNA during cell lysis, bacterial RNA, and co-isolated proteins, complicating their effective isolation, sequencing, expression quantification and downstream analyses (Ramirez, et al., 2018).

Standard RNA-seq bioinformatics pipelines, predominantly designed for abundant and high-quality cellular RNAs, inadequately handle the extremely low abundance, fragmented nature, and contamination-prone characteristics of EV RNAs. For instance, conventional poly-A selection sequencing preferentially enriches protein-coding transcripts, neglecting crucial non-polyadenylated RNAs abundant in EVs, such as miRNAs and lncRNAs. Furthermore, traditional approaches typically fail to adequately manage contamination risks and multiple mapping reads, potentially resulting in false-positive findings, missed valuable RNA species and reduced reliability. Therefore, specialized bioinformatics pipelines tailored to EV total RNA-seq are urgently required for accurate annotation, quantification, and interpretation of complex EV RNA profiles.

We systematically reviewed methodologies for EV isolation, RNA extraction, and library preparation, evaluating their strengths and limitations. Traditional EV isolation methods, such as ultracentrifugation, generally exhibit lower efficiency and higher variability. In contrast, newer approaches such as the Exodos system offer improved purity, size uniformity (30-200 nm), efficiency and reproducibility. For RNA extraction (Chen, et al., 2021), the optimized TRIzol method is advantageous for EV samples characterized by low RNA yield and fragmented RNA, effectively preserving RNA integrity and minimizing DNA contamination (Mateescu, et al., 2017; Prendergast, et al., 2018). Regarding RNA-seq library construction, we recommend the SMARTer Pico v3 kit, given its superior sensitivity, robustness, and compatibility with ultra-low-input RNA samples at picogram levels typical of EV preparations. This kit effectively captures diverse RNA species, including short, fragmented, and non-polyadenylated RNAs by leveraging strand-specific RNA library construction and unique molecular identifier (UMI) technologies, enabling comprehensive profiling of heterogeneous EV RNA populations with minimal bias and contamination. Detailed methodological comparisons are provided in the supplementary materials.

To address these analytical challenges (Miceli, et al., 2024), we developed EVscope, an end-to-end computational pipeline designed to automate and standardize EV total RNA-seq analyses. EVscope incorporates a novel genome-wide EM-based algorithm for quantifying multi-mapping reads and integrates comprehensive quality control measures, RNA classification, and downstream analytical components, including the inference of the cellular and tissues origins of EV RNAs. This comprehensive workflow significantly enhances the reproducibility, robustness and scalability of EV RNA-seq analyses, providing an efficient tool for investigating EV-derived RNA signatures.

## 2. Materials and methods

EVscope is designed as a fully automated pipeline that processes raw RNA-seq data from EV samples and generates a comprehensive set of quality control reports, expression data, and visualization outputs. Detailed comparisons of EV isolation methods (EXODUS system), RNA extraction efficiency (TRIzol-based vs. column-based methods), RNA library preparation (SMARTer Pico v3 kit), and quality control assessments (adapter trimming and read length distributions) are provided in Supplementary Figures S1-S4. EVscope performance was rigorously validated using publicly available EV RNA-seq datasets (Table S1), highlighting the pipeline’s capability of accurately handling adapter trimming and multi-mapping read assignment. All analysis parameters and software versions are documented in the reproducible Conda environment provided with the pipeline. The pipeline is structured into the following major steps:

**Figure 1:**
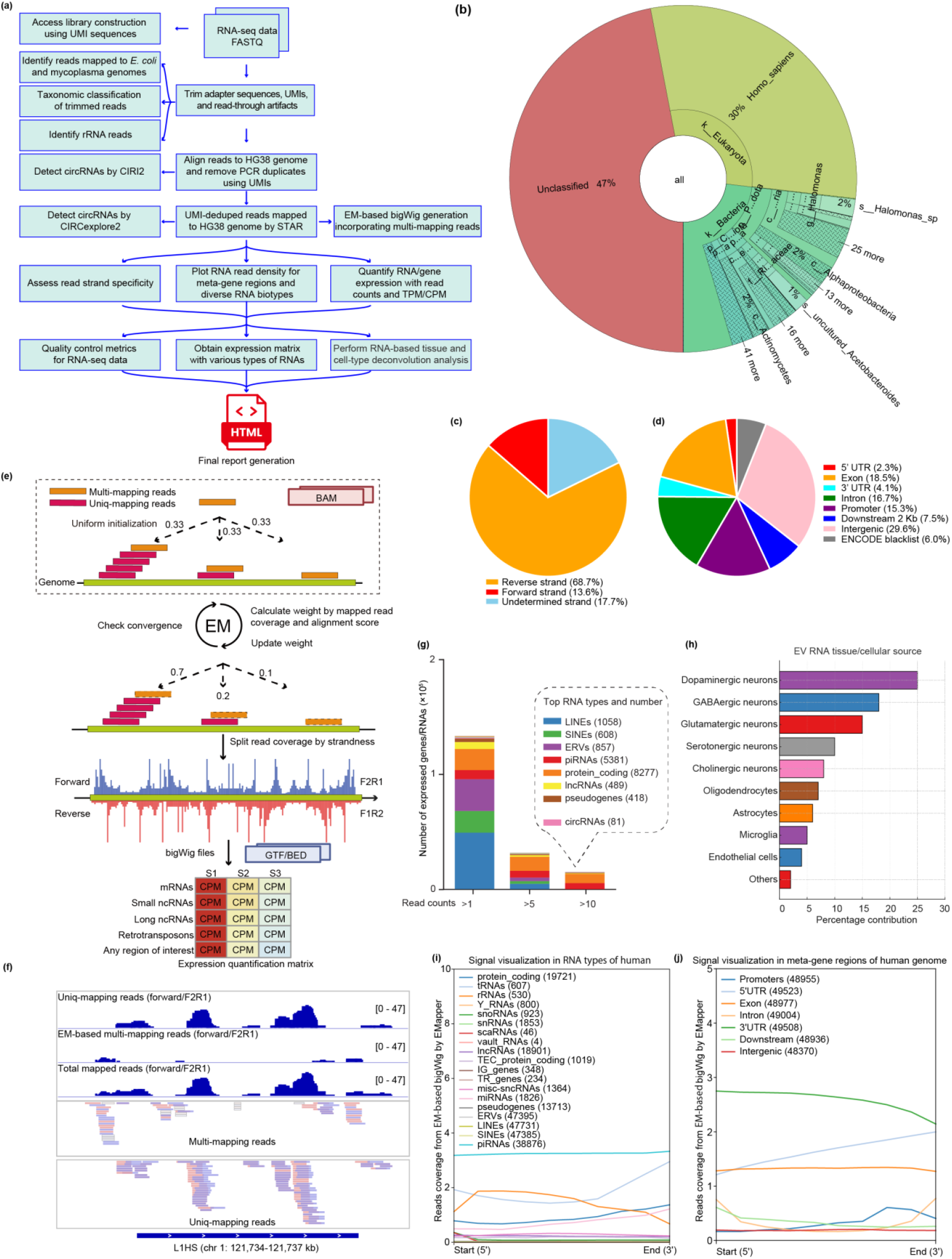
EVscope: A Comprehensive Bioinformatics Pipeline for Accurate and Robust Analysis of Total RNA Sequencing from Extracellular Vesicles. (a) Schematic overview of the EVscope bioinformatics workflow, highlighting key steps including quality control, contamination assessment, technical sequence trimming, read alignment, RNA quantification, EM-based BigWig generation and downstream analyses. (b) Taxonomic classification and distribution of EV RNA-seq reads using Kraken, visualized by a Krona plot highlighting predominant contaminants (e.g., bacteria) and unclassified reads. (c) Strand-specific analysis showing the distribution of read strand specificity. (d) Proportions of RNA-seq reads mapped to distinct genomic regions, represented by read count percentages. (e) Illustration of the Expectation-Maximization (EM) algorithm-based approach used by EVscope for accurate quantification and assignment of multi-mapped reads at single-base resolution, including a detailed example of BigWig tracks comparing uniquely mapped and EM-corrected multi-mapped reads. (f) Integrative Genomics Viewer (IGV) visualization example of EM algorithm-corrected BigWig tracks demonstrating enhanced resolution for multi-mapping read assignments. (g) Bar plot depicting the distribution and number of expressed RNAs across distinct biotypes within EV RNA-seq data. (h) Predicted cellular and tissue origins of EV-derived RNAs using single-cell RNA sequencing (scRNA-seq) deconvolution, displaying cell-type contributions ranked by abundance. (i) Visualization of EV RNA-seq signals across RNA biotypes in the human genome, highlighting biotype-specific EM-based read coverage patterns generated by EMapper. (j) Visualization of EV RNA-seq signals within meta-gene regions across the human genome, illustrating EM-based read coverage distribution across annotated genomic features by EMapper.

### Quality control (QC) and UMI motif visualization

Quality control of both raw sequencing reads and final cleaned reads after UMI-adapter trimming was performed using FastQC (https://www.bioinformatics.babraham.ac.uk/projects/fastqc) to ensure high sequencing data quality for downstream analyses. Unique Molecular Identifier (UMI) motif composition was visualized using custom Python scripts and inspected for correctness and structural integrity. For human EV RNA-seq libraries, we recommend using the SMARTer® Stranded Total RNA-Seq Kit v3 – Pico Input Mammalian (Takara Bio), where the first 14 nucleotides of Read2 contain a structured UMI sequence consisting of a UMI-linker-UMI-adapter motif, which includes adapter and linker sequences. Specifically, the constant presence and integrity of the UMI linker sequences within the reads were confirmed visually as part of high-quality validation for RNA-seq library construction.

### Read Contamination Assessment

Bacterial contamination from human sources is detected and filtered using BBMap (v39.15) (https://sourceforge.net/projects/bbmap), which partitions reads against bacterial reference genomes, including *Escherichia coli* and Mycoplasma species (NCBI genome: https://www.ncbi.nlm.nih.gov/home/genomes). Kraken2 (Wood, et al., 2019) is used for taxonomic classification of reads, providing an overview of microbial and viral contaminants. The results are visualized using Krona (Ondov, et al., 2011) interactive plots. To address ribosomal RNA (rRNA), which can constitute a major fraction of total RNA libraries, EVscope uses RiboDetector (Deng, et al., 2022) on downsampled reads to accurately quantify rRNA contamination, enhancing assessment precision.

### UMI processing, adapter and UMI-read through trimming

A multi-stage trimming process is implemented to prepare reads for alignment. First, Unique Molecular Identifiers (UMIs) are extracted from Read2 using UMI-tools (Smith, et al., 2017) and appended to the read headers for subsequent deduplication. Second, standard sequencing adapters are trimmed using cutadapt (Martin, 2011). Third, a novel module (UMIAdapterTrimR1.py) specifically addresses the potential for UMI “read-through” artifacts, which can occur in short-fragment libraries by trimming UMI sequences that have been erroneously incorporated into the 3’ end of Read1. Finally, reads undergo quality trimming with cutadapt to remove low-quality bases, ensuring only high-quality reads proceed to alignment.

### Read strandedness detection

The strandedness specificity of the RNA-seq library is determined using a customized RSeQC-based script (Wang, et al., 2012). Leveraging this strandedness information, EVscope optionally quantifies reads from both RNA-derived (correct strand) and potential genomic DNA-derived (opposite strand) sources separately using featureCounts (Liao, et al., 2013). Subsequently, the read counts from the incorrect strand are then subtracted from the correct strand’s counts to produce a gDNA-corrected expression profile, a critical step for accurately analyzing low-input total RNA samples. RNA integrity and other library metrics are collected using Picard tools (https://broadinstitute.github.io/picard).

### Alignment and various types of linear RNA identifications

Reads are aligned to the human reference genome (hg38) using STAR (Dobin, et al., 2012) in a two-pass mode with parameters (e.g., --outFilterMultimapNmax 100, --chimSegmentMin 10) to enhance mapping accuracy for EV-derived RNA fragments. RNA species are annotated using a comprehensive set of annotation references. The primary gene annotation is based on GENCODE comprehensive v45, which is supplemented with piRNA annotations from piRBase (Wang, et al., 2022) and retrotransposon repetitive element annotations from UCSC genome browser by RepeatMasker (Chen, 2004) and Dfam (Hubley, et al., 2016). This integrated annotation allows for the classification and quantification of a wide array of RNA biotypes, including protein-coding genes, circRNAs, piRNAs, LINEs, SINEs, ERVs, and other non-coding RNAs. Read counts are assigned to genomic features using either featureCounts (Liao, et al., 2013) for uniquely mapping reads or optionally RSEM (Li and Dewey, 2011) for multi-mapping read quantification. Expression matrices are generated in TPM and CPM formats.

### Circular RNA identification

Circular RNA identification was performed using two complementary computational methods, CIRI2 and CIRCexplorer2, to balance sensitivity and specificity while reducing false positives. CIRI2 (Gao, et al., 2017) was utilized first, as it excels in identifying novel circRNAs without reliance on prior annotations, offering a balanced performance between sensitivity and precision. Subsequently, circRNA candidates were further refined using CIRCexplorer2 (Zhang, et al., 2016), a method leveraging the comprehensive human GENCODE annotations v45 (Harrow, et al., 2012) (https://www.gencodegenes.org/human) to enhance the annotation accuracy, reduce false positives, and improve confidence in circRNA characterization.

### Single-base resolution read coverage generation using expectation-maximization (EM) algorithm for multi-mapping read assignment

We developed an expectation-maximization (EM) algorithm-based approach to accurately assign multi-mapped sequencing reads at single-base resolution, iteratively refining fractional alignments based on alignment scores (AS) and local read coverage derived from both uniquely and multi-mapped reads until convergence. We developed a novel tool, which we named EMapper, to implement this expectation-maximization (EM) algorithm. The EM algorithm (EMapper) initializes the fractional assignments uniformly across all potential alignments and iteratively refines them based on reads coverages and alignment scores (AS) until convergence, producing strand-specific, single-base resolution read BigWig coverage files. Expression quantification is performed by calculating the Mean Per-base CPM (MCPM), providing robust normalization suitable for cross-sample and gene-level comparisons and downstream analyses such as differential expression and functional association studies.

### Expression quantification and RNA type characterization

EVscope generates RNA distribution plots and expression matrices, providing an overview of RNA biotype composition. BigWig tracks are generated to visualize RNA coverage for both uniquely mapped reads and multi-mapped reads corrected by an Expectation-Maximization (EM) algorithm.

### Cellular origin inference

To determine the likely source of EV RNAs, EVscope integrates the ARIC deconvolution algorithm (Zhang, et al., 2022), which robustly estimates cell-type proportions from bulk RNA-seq data through systematic marker selection. Users can input custom bulk or single-cell reference datasets or leverage built-in references with GTEx v10 (Lonsdale, et al., 2013) and the Human Brain Cell Atlas v1.0 (Siletti, et al., 2023) to infer the cellular origins of EV-derived RNAs.

### HTML final report generation

The final HTML report for EV RNA-seq analysis using EVscope was generated using R Markdown (Xie, et al., 2018). Interactive data visualization and tabular summaries were prepared using the DT package (https://rstudio.github.io/DT/), and formatted tables were generated using the kableExtra package (Zhu, 2021). The overall document structure and navigation were organized with the bookdown package (Xie, 2016), providing clear sectioning and interactive table-of-contents navigation. Project paths and reproducibility were managed by the here package (Müller, 2020). The final report includes tabbed sections, interactive tables allowing direct export of data, and embedded external reports (e.g., QC reports from FastQC and cutadapt).

## 3. Conclusion

EVscope has been tested on multiple EV RNA-seq datasets, demonstrating effective contamination filtering, RNA classification, and cellular origin inference. EVscope systematically generates 27 structured output directories corresponding to each major pipeline step, providing detailed and clearly organized results, including QC reports, expression data, BigWig coverage files, and interactive HTML reports. EVscope is implemented as a Bash script, allowing easy execution and custom modifications. The full source code and user documentation are available at GitHub (https://github.com/TheDongLab/EVscope) and archived on Zenodo (https://zenodo.org/records/15577789) under an MIT open-source license. EVscope provides an automated, scalable solution for analyzing total RNA-seq data from extracellular vesicles. By incorporating comprehensive quality control, contamination detection, broad RNA biotype classification, and cellular origin inference, EVscope addresses key challenges in EV RNA-seq research. This tool standardizes data processing workflows, enhances reproducibility, and facilitates the discovery of EV RNA biomarkers.

## Author contributions

Yiyong Zhao (Conceptualization [supporting], Data curation [lead], Formal analysis [lead], Software [lead], Visualization [lead]), Himanshu Chintalapudi (Visualization [supporting]), Ziqian Xu (Resources [supporting]), Weiqiang Liu (Data curation [supporting]), Yuxuan Hu (Validation [supporting]), Ewa Grassin (Resources [supporting]), Minsun Song (Resources [supporting]), SoonGweon Hong (Resources [supporting]), Luke P. Lee (Resources [supporting]) and Xianjun Dong (Conceptualization [lead], Methodology [supporting], Funding acquisition [lead], Project administration [lead], Supervision [lead]).

## Supplementary data

Supplementary data are available at Bioinformatics online.

## Data availability

Source code is available at GitHub (https://github.com/TheDongLab/EVscope) and archived on Zenodo (https://zenodo.org/records/15577789) under an MIT open-source license. Raw sequencing data used for validation are publicly available from NCBI Sequence Read Archive (SRA accession: SRR31350808-11). For any inquiries, please contact Dr. Xianjun Dong at.

## Supplementary Material and methods

### Expectation-maximization (EM) algorithm for multi-mapping read assignment at single-base resolution

In our EM algorithm, a multi-mapped fragment or read *r* (whether paired-end or single-end) can align to multiple genomic loci. For a given fragment/read *r*, let its alignments be represented as:

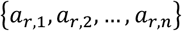

The alignment score (AS) in the BAM file reflects mapping quality determined by alignment tools (e.g., STAR, Hisat2, BWA). Typically, these scores—based on nucleotide matches, mismatches, and gap penalties—are stored in tags such as the “AS:i” tag used by STAR. We denote the AS scores for alignments of fragment/read *r* as:

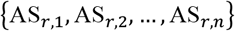

Initially, we assume that a multi-mapped fragment/read r originates uniformly from any of its potential source loci. Consequently, each alignment is assigned an equal fractional weight of:

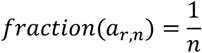

This uniform initialization treats all candidate loci equally, simplifying initial computations and providing an unbiased starting point. This approach enables the EM algorithm to robustly converge through iterative refinement based on observed coverage and alignment quality. Each alignment *a*_*r,n*_ covers a genomic interval from start_*r,n*_ to end_*r,n*_. At a given genomic position *p*, the coverage contributed by uniquely mapped reads is represented as Cov^uniq^(*p*). At iteration *k*, the weight for each alignment *n* is computed as:

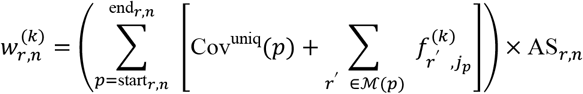

Where:

ℳ(*p*) is the set of multi-mapped reads covering genomic position *p*.

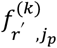 is the fractional assignment (weight) of multi-mapped read *r*^′^ at alignment *j*_*p*_ specifically covering genomic position *p* at iteration *k*.

These weights are converted into the fractional contribution for the next iteration:

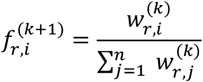

After updating fractions for all multi-mapped reads, the coverage arrays are recalculated by aggregating each fractional read contribution across corresponding genomic intervals. This iterative process continues until the total fractional change between successive iterations falls below a user-defined tolerance *ε*, or a predefined maximum number of iterations is reached. The convergence criterion is formally defined as:

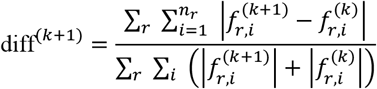

When diff^(*k*+1)^ < *ε*, convergence is declared. The final fractional assignments for multi-mapped reads are then incorporated into the coverage arrays, which can subsequently be used for downstream analyses, such as gene expression estimation. In our implementation, we recommend setting the convergence threshold to *ε* < 1e^-3^ or limiting the maximum number of iterations to 200. The algorithm reliably converges and generates EM-based single-base resolution read coverage files with strand information in BigWig format for further targeted gene or region expression quantification, as well as for further analyses, including the exploration of expression associations with neighboring regions.

### Measurement of expression quantification for gene/interested region from EM-based BigWigs at single-base resolution

We provide continuous, single-base resolution BigWig files with strand information. Users can run bigWig2CPM.py to calculate the mean per-base CPM (MCPM) for any genomic region, enabling cross sample expression comparisons and facilitating downstream analyses such as differential gene expression analysis and expression association studies with host RNA or neighboring regions for detection of dependent or independent actively transcribed elements. Counts per million (CPM) is a normalization approach that scales coverage or read counts by the total number of reads, in millions, to account for differences in library size.

We define the per-base CPM as:

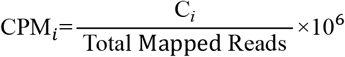

where C_*i*_ is the coverage at base *i*, then the MCPM over a gene/region of length *L* is then given by:

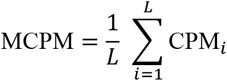

This MCPM value reflects the average normalized coverage across the region, facilitating robust expression quantification and cross sample comparisons.

### Comparison and recommendation of EV isolation

Here, we recommend EXODUS, an automated EV (extracellular vesicle) isolation system. Exosomes, typically ranging from 30 to 200 nm, represent a major subset of EVs (Chen, et al., 2021). The isolation process involves filtering the raw sample-such as plasma, cell culture medium, urine, or tears—either after appropriate dilution or directly without dilution. For example, to isolate EVs from human plasma, the plasma should first be centrifuged at 12,000 × g for 30 min at 4°C. Then, 400 µL of the supernatant is diluted in 30 mL of PBS. The sample is then filtered through a 0.22 µm filter using a syringe before processing on the instrument. The filtered sample is then placed into the system’s sample reservoir, and an exosome isolation device (EID) chip compatible with the experimental requirements is selected. Upon running the corresponding program and inputting the necessary parameters, such as sample name, sample type, input volume, and tube capacity, the system automatically isolates exosomes from the sample.

**Figure S1.**
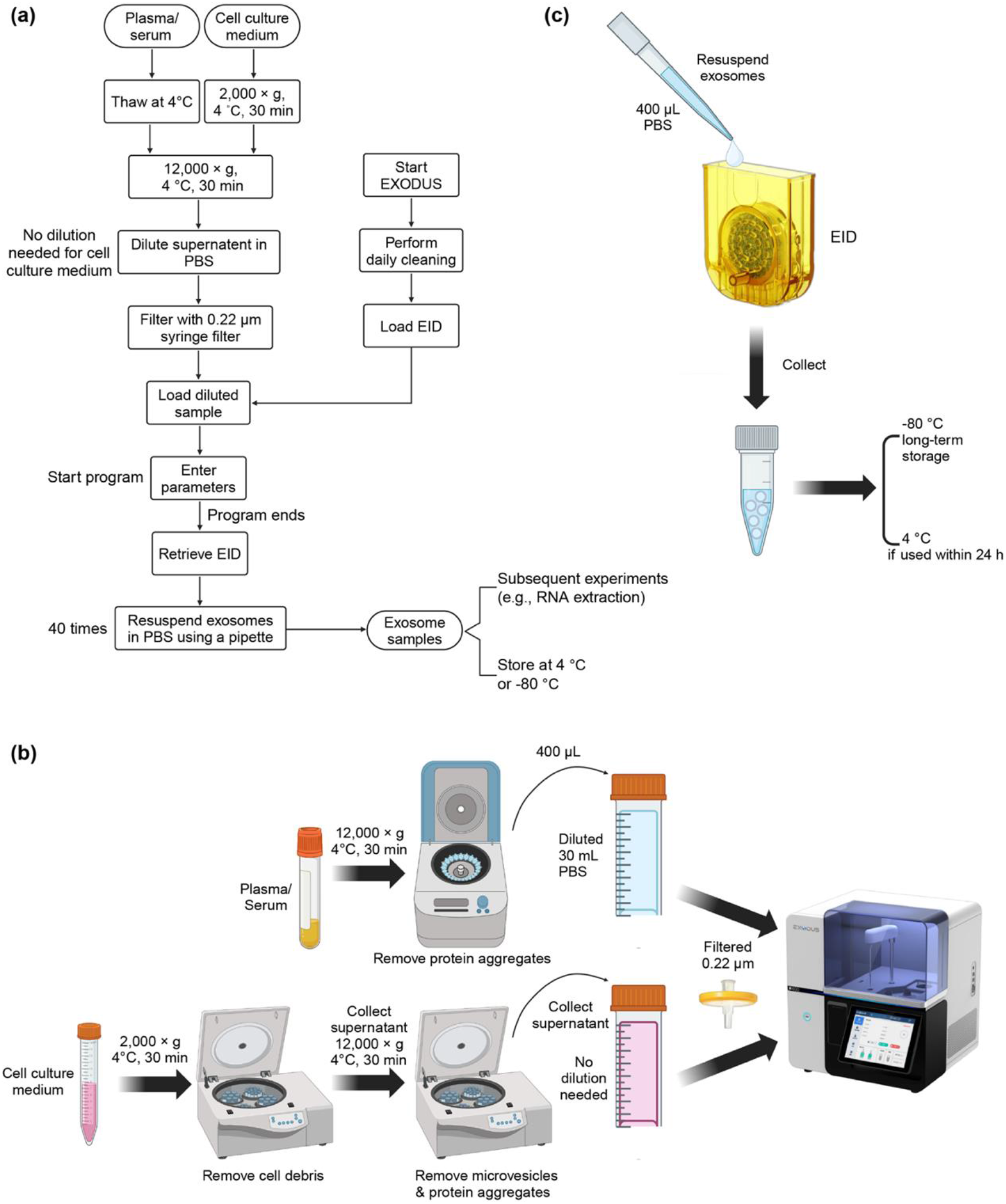
Exosome isolation workflow using the recommended EXODUS method (Chen, et al., 2021). **(a)** Schematic workflow for isolating exosome from plasma/serum or cell culture medium via the EXODUS platform (Chen, et al., 2021). **(b)** Detailed pre-processing steps required for plasma/serum and cell culture medium samples before loading into the EXODUS system (Chen, et al., 2021). **(c)** Illustration of the exosome resuspension step from the Exosome Isolation Device (EID).

The core structure of the EID chip consists of a chamber enclosed by two nanoporous membranes. The raw sample is introduced into the chamber in a stepwise manner. The system, connected to the exterior of the membranes via tubing, generates ultrasonic oscillations, driving the sample to pass through the membranes from both sides. Particles larger than the membrane pore size, such as exosomes, are retained on the membrane surface within the chamber. The membrane pore size is approximately 30 nm. Since the sample is filtered through a 0.22 µm filter before processing, the final particles obtained should theoretically range from 30 to 220 nm in diameter, which aligns with the typical size of exosomes. After all sample fractions have passed through the membranes, the chamber is rinsed to resuspend and recover the EVs adhered to the membrane.

### Comparison and recommendation of EV RNA extraction

We employed three different methods to isolate RNA from exosomes, including two commercial column-based extraction kits and the traditional TRIzol precipitation method. Not all methods are suitable for the extraction of exosomal RNA. Due to the typically low abundance and small fragment size of exosomal RNA, our experimental comparisons suggest that silica column-based methods are not optimal for exosomal RNA extraction.

We evaluated the performance of two commercial extraction kits, miRNeasy Micro Kit (QIAGEN, catalogue number: 217084) and miRNeasy Serum/Plasma Advanced Kit (QIAGEN, catalogue number: 217204). The primary difference between the two kits lies in the composition of the lysis buffer; one contains phenol, while the other is phenol-free, offering reduced toxicity. Both methods operate on the principle of sample lysis followed by phase separation. The RNA-containing supernatant is applied to a silica column, where RNA binds to the silica, followed by multiple rounds of washing. Finally, RNA is eluted by RNase-free water. To ensure RNA purity, we performed on-column DNase I treatment (QIAGEN, catalogue number: 79254).

Total RNA was extracted using the QIAGEN miRNeasy Micro Kit following the manufacturer’s protocol. We consider exosome samples as low-concentration plasma or cell samples. Briefly, 700 µL QIAzol Lysis Reagent was added to the sample, vortexed, and incubated at room temperature (15-25 °C) for 5 min. After the addition of 140 µL chloroform, the mixture was shaken vigorously for 15 s, incubated for 2-3 min, and centrifuged at 12,000 × g for 15 min at 4°C to achieve phase separation. The upper aqueous phase was carefully transferred to a new tube, mixed with 1.5 × volumes of 100% ethanol, and loaded onto a RNeasy MinElute spin column. After centrifugation at 8000 × g for 15s, the flow-through was discarded, and the remaining sample was processed similarly. DNase I treatment was performed by adding 80 µL of DNase I solution (10 µL DNase I stock and 70 µL Buffer RDD) directly onto the spin column membrane, followed by a 15-min incubation at 20-30 °C. The membrane was then washed sequentially with Buffer RWT, Buffer RPE, and 80% ethanol, with centrifugation steps at 8000 × g. To thoroughly removal of residual ethanol, the column was centrifuged at full speed for 5 min before RNA elution. RNA was eluted using 18 µL RNase-free water, quantified with a NanoDrop, and stored at - 80 °C for downstream analysis.

Total RNA extraction from exosome samples using the QIAGEN miRNeasy Serum/Plasma Advanced Kit is also following the manufacturer’s instructions. Briefly, 200 µL of exosome sample was mixed with 60 µL Buffer RPL, vortexed for 5s, and incubated at room temperature (15-25 °C) for 3 min. After adding 20 µL Buffer RPP, the lysate was vortexed for 20 s to ensure thorough mixing and incubated for 3 min. Phase separation was achieved by centrifugation at 12,000 × g for 3 min at room temperature, and the resulting clear supernatant (230 µL) was transferred to a new tube containing an equal volume of isopropanol. The mixture was loaded onto a RNeasy UCP MinElute spin column, centrifuged at 8000 × g for 15s, and the flow-through was discarded. The same DNase I treatment was performed by adding 80 µL of DNase I solution directly to the membrane, followed by a 15-min incubation at room temperature. The membrane was washed sequentially with Buffer RWT, Buffer RPE, and 80% ethanol, with centrifugation steps at 8000 × g. The spin column was then dried by centrifugation at 12,000 × g for 5 min. RNA was eluted using 18 µL RNase-free water and quantified using a NanoDrop.

**Figure S2.**
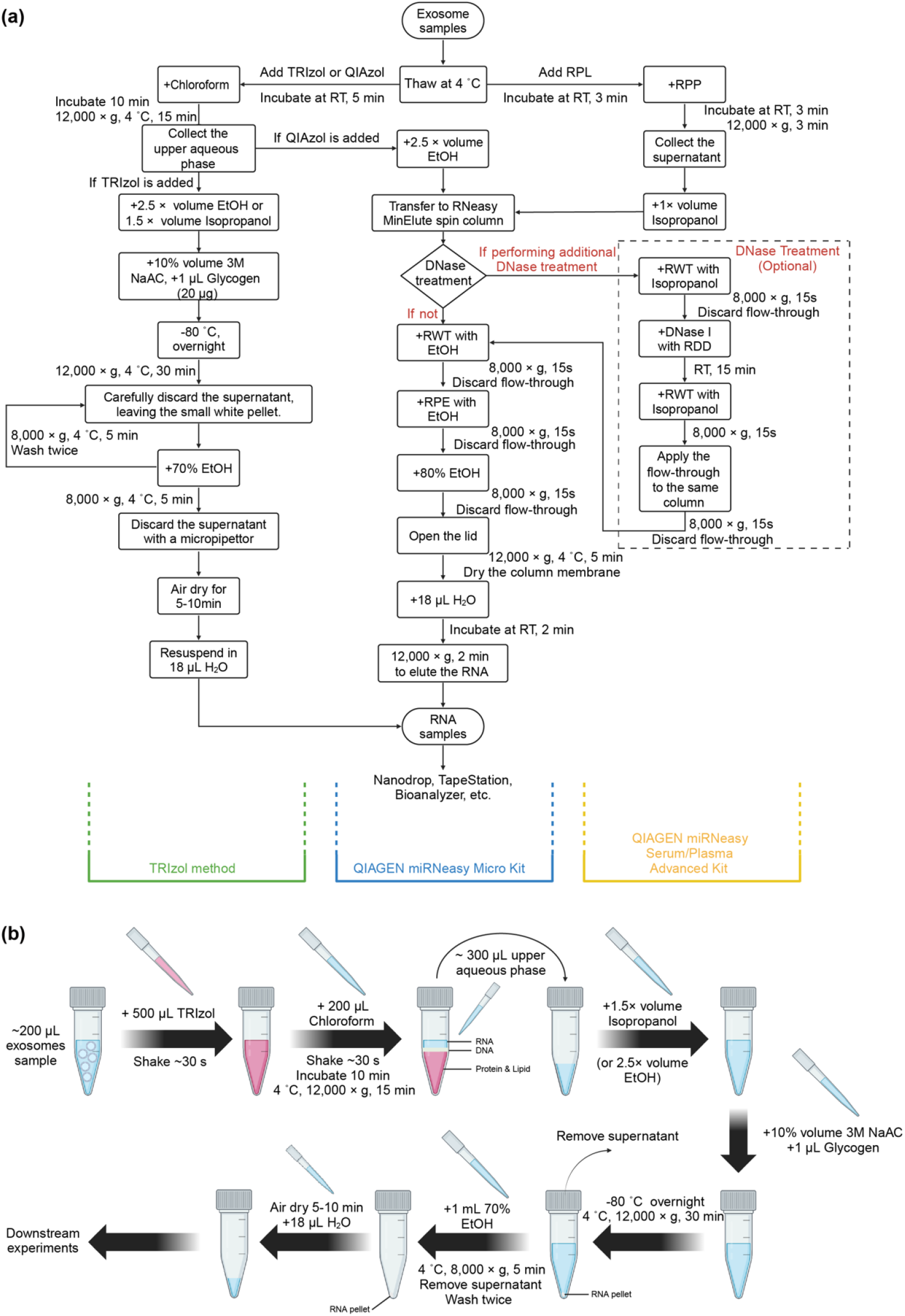
Conventional recommended column/solution-based workflows for exosomal RNA extraction. **(a)** Workflow comparison of three exosomal RNA isolation methods: two commercial QIAGEN (miRNeasy Micro Kit and miRNeasy Serum/Plasma Advanced Kit and an optimized TRIzol-based method (Tang, et al., 2017; Xu, et al., 2018). **(b)** Schematic overview illustrating detailed steps of the optimized TRIzol-based method exosomal RNA extraction (Prendergast, et al., 2018).

We highlight an improved TRIzol-based extraction method, highly recommended for EV RNA extraction (Prendergast, et al., 2018). The traditional TRIzol method for RNA precipitation appears to be more suitable for exosomal RNA extraction. During the experiment, the formation of a clear precipitate allowed for direct assessment of RNA extraction efficiency, providing an immediate indication of successful RNA isolation. Briefly, 200 µL of isolated exosomes (containing at least 10^^^8 total exosomes) were mixed with 500 µL TRIzol reagent (Invitrogen, catalogue number: 15596026) by shaking, followed by incubation on ice for 5 min to ensure complete dissociation of nucleoprotein complexes. Subsequently, 200 µL of pure chloroform was added, mixed thoroughly for 30 s, and incubated for 3 min. Phase separation was achieved by centrifugation at 12,000 × g for 15 min at 4°C, resulting in the formation of a lower red phenol-chloroform phase, an interphase, and an upper colorless aqueous phase. The RNA-containing aqueous phase was carefully transferred to a new collection tube, avoiding contamination from the interphase. RNA was precipitated by adding 1.5 × volume of isopropanol (or alternatively 2.5 × volume of 100% ethanol), 10% volume of sodium acetate [3M, pH 5.5] (Invitrogen, catalogue number: AM9740), and 1 µL ultrapure glycogen (20 µg/µL) (Invitrogen, catalogue number: 10-814-010), followed by overnight incubation at -80 °C to ensure complete RNA precipitation. The next day, the sample was centrifuged at 12,000 × g for 30 min at 4°C to pellet the RNA. The pellet was washed twice with 1 ml of 70% ethanol, centrifuged at 8,000 × g for 5 min at 4°C, and air-dried for 20 min at room temperature. Finally, the RNA was resuspended in 18 µL of RNase-free water for further analysis.

We compared the RNA extracted by the three methods using NanoDrop results. For the first two kit-based methods, the NanoDrop spectra showed no detectable peaks, indicating that RNA was not successfully extracted or was below the detection limit. In contrast, for the TRIzol method, a distinct absorption peak was observed at 260 nm, with a trough at 230 nm, suggesting that this method allows us to extract pure and high-quality exosomal RNA. In addition, we also assessed RNA quality using the high-sensitivity RNA TapeStation assay. The results indicated that for exosomal RNA extracted using the miRNeasy Micro Kit, no bands or peaks were observed, regardless of whether DNase I treatment was applied, suggesting that no significant amount of RNA was extracted. In contrast, for RNA extracted using the miRNeasy Serum/Plasma Advanced Kit, DNase I treatment again failed to produce any bands or peaks, whereas no additional DNase I treatment allowed bands and peaks to be observed in several samples. For the improved TRIzol method, bands were always visible. It is important to note that due to the low content of exosomal RNA, which typically lacks 18S and 28S rRNA, RIN values are generally not expected on the TapeStation.

### Comparison and recommendation of RNA Library preparation for EV RNA sequencing

Efficient library preparation is critical for accurate RNA sequencing of extracellular vesicles (EVs). Here, we compared several commercially available RNA library preparation kits: SMARTer Pico v3, SMARTer HI, SMARTer HT, RNA Access, RNA Exome, KAPA Hyper, Ovation SoLo, and KAPA Hyper UMI. SMARTer Pico v3 is optimized specifically for low-input RNA, making it particularly suitable for EV-derived RNA, which is typically present at low abundance and variable quality. The kit exhibits high sensitivity, improved accuracy, and minimal amplification bias, enabling robust detection of rare transcripts (Hagemann-Jensen et al., 2020; Srinivasan et al., 2019). In contrast, SMARTer HI and HT kits are tailored for high-throughput applications but are less effective for ultra-low input RNA and show increased bias at low input concentrations (Mereu et al., 2020). RNA Access and RNA Exome kits are designed primarily for targeted sequencing and perform poorly for unbiased transcriptome profiling, limiting their suitability for comprehensive EV RNA sequencing (Archer et al., 2014). KAPA Hyper and KAPA Hyper UMI kits excel in high-complexity RNA samples and provide unique molecular identifiers (UMIs) to reduce PCR duplication bias, but their efficiency markedly declines at extremely low RNA inputs typical of EV samples (Ziegenhain et al., 2017). Similarly, Ovation SoLo, while optimized for low RNA input, demonstrates inconsistent performance and variable coverage efficiency for EV RNA (Leinonen et al., 2017). Given these considerations, the SMARTer Pico v3 emerges as the most suitable choice for EV RNA sequencing, owing to its superior performance with low input, sensitivity, reproducibility, and transcriptomic fidelity.

Given the extremely low abundance and trace quantities of RNA within extracellular vesicles (EVs), reliable RNA-sequencing (RNA-seq) library preparation remains technically challenging. Conventional library construction methods, such as standard mRNA or total RNA sequencing protocols, frequently lack sufficient sensitivity and robustness for EV-derived RNA. These methods often lead to significant RNA loss or degradation due to multiple enzymatic treatments, purification steps, and inefficient adapter ligation, thereby compromising the reliability and reproducibility of downstream analyses.

Previously employed RNA-seq protocols for EVs include traditional poly(A)-tail enrichment methods and ribosomal RNA (rRNA) depletion approaches. Poly(A)-tail enrichment methods inherently bias against non-polyadenylated RNAs, leading to incomplete transcriptomic profiles and underrepresentation of important RNA classes such as non-coding RNAs prevalent in EVs (Huang, et al., 2013; Van Balkom, et al., 2015). Conversely, rRNA depletion methods, while preserving non-polyadenylated transcripts, still frequently result in substantial RNA loss during depletion steps, which negatively impacts sensitivity and detection accuracy, particularly in low-input scenarios like EV RNA (Enderle, et al., 2015; Mateescu, et al., 2017). Furthermore, methods involving standard adapter ligation protocols have historically been inefficient in capturing short RNA fragments or degraded RNAs, which are abundant in EV preparations, further limiting their applicability and accuracy (Srinivasan, et al., 2019).

Recent advancements in library preparation techniques, particularly the SMARTer Stranded Total RNA-Seq Kit v3 – Pico Input Mammalian (Takara Bio), have provided substantial improvements tailored explicitly for extremely low-input RNA samples, such as those obtained from EVs. This strategy employs an efficient template-switching mechanism coupled with unique molecular identifiers (UMIs), enabling accurate amplification, quantification, and normalization of scarce RNA molecules. Additionally, the stranded nature of this library preparation retains critical information regarding RNA strand orientation, essential for detailed transcriptomic analysis and accurate identification of antisense transcripts. Recent literature has consistently demonstrated that this library preparation strategy delivers superior sensitivity, accuracy, and reproducibility when applied to EV-derived RNA, effectively overcoming previous limitations associated with RNA yield and integrity (Neri, et al., 2022; Shi, et al., 2021). Based on current RNA-seq methodological advancements and empirical evidence in published research, we strongly recommend employing the SMARTer Stranded Total RNA-Seq Kit v3-Pico Input Mammalian for EV RNA-seq library construction. This approach provides distinct advantages in sensitivity, reproducibility, strand specificity, and quantitative accuracy, making it ideally suited for the reliable characterization and functional analysis of EV transcriptomes. Furthermore, within the EVscope platform, we have integrated a specialized module that directly visualizes the UMI region’s nucleotide composition and base density from raw sequencing reads, thereby providing immediate quality assessment and validation of library construction efficacy.

### Illumina adapters and UMIs trimming and UMI-based deduplication of paired-end reads

Adapter trimming and deduplication of paired-end sequencing reads were performed using Illumina and UMI-based technologies. To accurately quantify RNA expression, particularly small RNAs derived from total RNA sequencing, obtaining high-quality, adapter-free sequencing reads is critical. Paired-end reads generated by the SMARTer Stranded Total RNA-Seq Kit v3 contain unique molecular identifiers (UMIs) and associated adapter sequences that must be rigorously removed prior to downstream analysis. To ensure complete removal of technical adapter contamination, we employed a two-step trimming strategy. First, we extracted the initial 14 bp UMI sequences from the 5’ ends of Read2 using cutadapt. Due to the presence of short RNA inserts, read-through events occasionally occur, resulting in adapter contamination at the 3’ ends of Read1. Therefore, the extracted 14 bp UMIs from Read2 were reverse-complemented and subsequently used as adapter sequences to systematically trim the corresponding Read1 sequences. Adapter trimming was implemented using a custom Python script developed with Biopython (v1.81). The script performed 3’-end trimming of Read1 sequences, allowing a minimum overlap of 3 bp and a maximum mismatch rate of 10%. Reads shorter than 10 bp after adapter trimming were discarded from subsequent analyses. To efficiently process large-scale FASTQ datasets, we employed parallel computing using Python’s concurrent futures library, enabling simultaneous utilization of multiple CPU cores. Detailed quality control statistics, including the numbers of trimmed, short, and discarded reads, were recorded throughout the trimming process to ensure data integrity and to facilitate downstream RNA expression quantification.

**Figure S3.**
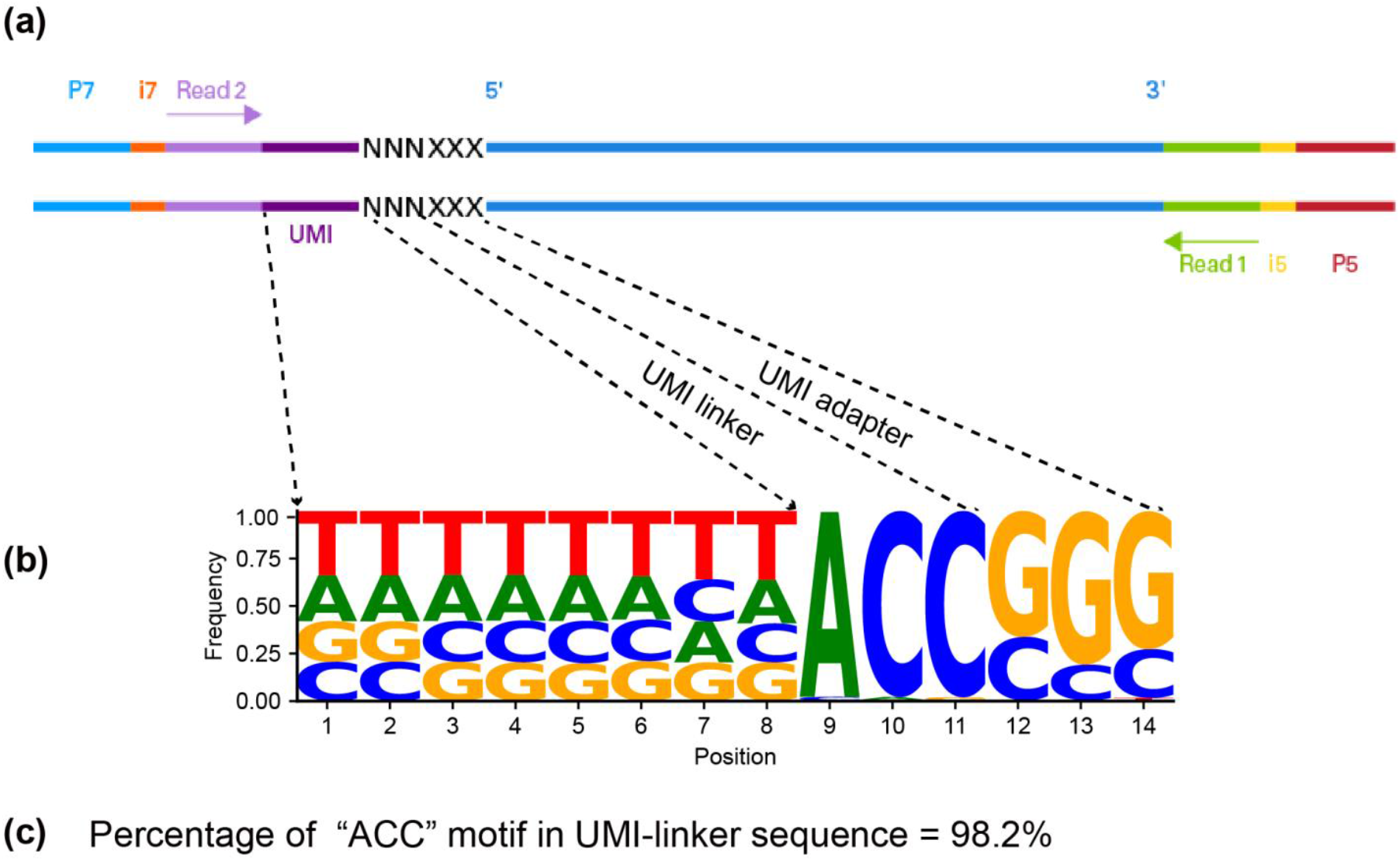
Structural features and UMI motif validation for EV RNA-seq libraries prepared using the SMARTer Stranded Total RNA-Seq Kit v3-Pico (Takara Bio). **(a)** Schematic representation of adapter sequences added via the SMARTer RNA Unique Dual Index Kit - 24U (Takara Bio, Cat. No. 634451). These adapters facilitate cluster generation on Illumina flow cells, with key regions highlighted: P7 (light blue), P5 (red), Index 1 (i7, orange), Index 2 (i5, yellow), sequencing primers Read Primer 2 (Read2, purple), and Read Primer 1 (Read1, green). Sequencing orientation is indicated relative to the original RNA (5′ and 3′, dark blue). The first 8 nucleotides of Read2 correspond to unique molecular identifiers (UMIs, dark purple), followed by a 3-nucleotide UMI linker (represented as NNN) and a 3-nucleotide sequence derived from the Pico v3 SMART UMI adapter (represented as XXX). **(b)** Sequence motif visualization of the first 14 nucleotides of Read2 (FASTQ format), representing the UMI region, was performed using EVscope’s “plot_fastq2UMI_motif.py” module. This QC step confirms successful library construction by displaying variable UMI sequences adjacent to a constant “ACC” UMI-linker and three fixed nucleotides from the adapter. This specific pattern confirms proper library preparation for EV RNA-seq using the Pico v3 SMART protocol. **(c)** The conserved “ACC” UMI-linker motif represented 98.2% of sequences analyzed, indicating high library construction efficiency (ideal: ∼100%). Takara Bio states that a fully successful RNA-seq library should exhibit a 100% “ACC” frequency. Thus, the observed percentage serves as an indicator of effective library construction. A significantly lower “ACC” frequency would suggest potential library preparation issues or sequencing errors. Additionally, the inclusion of a 10% PhiX spike-in is recommended for Illumina NovaSeq platforms by Takara Bio to enhance base diversity, sequencing quality, and accuracy in EV RNA-seq.

### Removal of UMI-derived technical sequences from Read1 via read-through

Raw sequencing reads (Read1) generated by the SMARTer-seq v3 kit were processed to eliminate technical sequences originating from unique molecular identifiers (UMIs). When the insert size of RNA or cDNA fragments is shorter than the sequencing read length, Read1 sequences inherently contain UMI-based technical sequences.

To specifically address this issue, we developed a custom Python pipeline (UMIAdapterTrimR1.py) designed to trim these sequences from Read1 by leveraging UMIs identified in the paired-end Read2. Initially, UMIs were extracted from the first 14 nucleotides (default setting: --umi-length=14) of Read2. Subsequently, reverse complement sequences of these UMIs were computed to define adapter sequences precisely. These adapters were then identified and trimmed from the 3’ ends of their corresponding Read1 sequences using a sliding window algorithm optimized for computational efficiency with Numba (Lam, et al., 2015). The trimming algorithm utilized a suffix-match approach, systematically evaluating potential overlaps from the full adapter length down to a specified minimum overlap of 3 nucleotides (−-min-overlap=3, default), allowing for a mismatch tolerance of up to 10% (−-error-rate=0.1, default). After trimming, reads shorter than 10 nucleotides (−-min-length=10, default) were discarded to maintain high-quality standards for downstream analyses. Formally, the adapter trimming procedure can be described as follows:

For each potential overlap length *L*, ranging from maximum adapter length down to the minimum overlap:

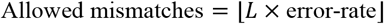

Where seq is the 3’-end of the read and adapter is the reverse-complemented UMI sequence. Mismatches between the sequence and the adapter are calculated as:

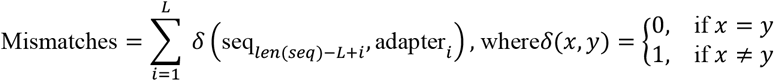

Trimming occurs when: mismatches ≤ allowed mismatches Detailed records of trimming events were logged in TSV format, capturing read identifiers, computed adapters, trimmed adapter sequences, and their lengths. Finally, cleaned Read1 sequences were output for subsequent analyses.

**Table S1:** Publicly available human EV RNA-seq datasets utilized for evaluating EVscope performance. EV RNA-seq datasets (NCBI SRA: SRR31350808-11) derived from neural stem cell EVs (LMNSC01 and LMNSC02) using the SMARTer Stranded Total RNA-Seq Kit v3-Pico (Takara Bio) and sequenced on an Illumina NextSeq 2000 platform. The table lists the NCBI SRA accession numbers, sample tissues, total sequencing yield, sequencing layout, instrument type, and publication dates. These samples served as benchmarks for validating EVscope’s adapter trimming accuracy, particularly its capability to identify and precisely remove UMI-derived adapter sequences from Read1, constructed by reverse complementing Read2 UMIs.

| NCBI SRA ID | Tissue | # of Bases | Layout | Instrument | Published |
| --- | --- | --- | --- | --- | --- |
| SRR31350808 | LMNSC02 neural stem cells rep2 | 14.7 Gb | Paired-end | NextSeq 2000 | 2024-11-21 |
| SRR31350809 | LMNSC02 neural stem cells rep1 | 16.7 Gb | Paired-end | NextSeq 2000 | 2024-11-21 |
| SRR31350810 | LMNSC01 neural stem cells rep2 | 17.4 Gb | Paired-end | NextSeq 2000 | 2024-11-21 |
| SRR31350811 | LMNSC01 neural stem cells rep1 | 12.7 Gb | Paired-end | NextSeq 2000 | 2024-11-21 |

EVscope was evaluated using four EV RNA-seq datasets detailed in Table S1, which were prepared using the SMARTer Stranded Total RNA-Seq Kit v3. Post trimming. Public datasets (Table S1) were specifically utilized to validate EVscope’s trimming efficiency, mapping accuracy, and multi-mapping resolution capability. We observed approximately 4.04-28.58% of Read1 sequences containing read-through technical sequences. Following application of our UMIAdapterTrimR1.py algorithm, the read length distributions of cleaned Read1 closely matched those of Read2. The removal of read-through technical sequences notably enhanced the detection of regulatory RNAs commonly enriched in EV samples.

**Figure S4.**
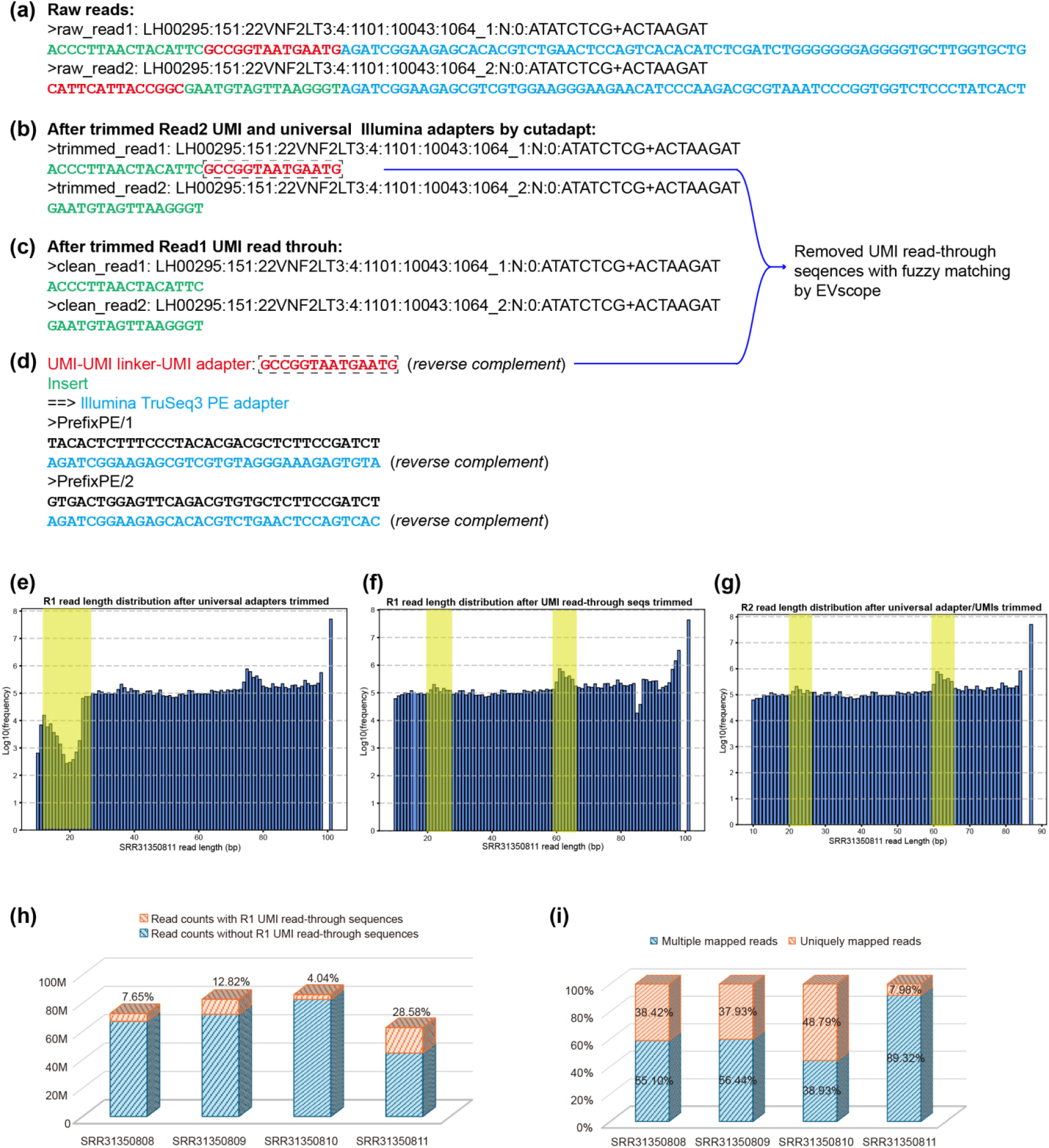
Comprehensive trimming and mapping analysis highlighting the necessity of the EM algorithm for multi-mapping read assignment in EV RNA-seq using EVscope. **(a)** Representative raw paired-end sequencing reads from EV RNA-seq libraries generated by the SMARTer Pico v3 protocol, highlighting the presence of technical UMI read-through sequences. (**b-c**) Sequential trimming workflow: Illumina universal adapters and UMI sequences were first trimmed using Cutadapt. Subsequent custom trimming using UMIAdapterTrimR1.py specifically addressed UMI read-through sequences frequently observed at the 3’ end of Read1, ensuring the accurate removal of technical artifacts. (d) Illustration of adapter sequences involved in library construction and trimming strategy. Specifically, it shows how EVscope constructs UMI-derived read-through adapter sequences utilizing reverse complements of Read2-derived UMIs to precisely remove artifacts from Read1 via fuzzy matching. (**e-g**) Read length distribution plots of a publicly available RNA-seq dataset processed through EVscope, demonstrating successive improvements in data quality. Panel (**e**) depicts read lengths after initial Illumina adapter trimming, revealing distinct length ranges strongly affected by residual UMI-derived read-through artifacts. Panels (**f-g**) present the distributions post-complete trimming using EVscope’s custom pipeline, demonstrating highly similar length distributions for cleaned Read1 and Read2. Highlighted regions correspond to regulatory RNA classes typically found in EVs: 20–30 bp (miRNAs, siRNAs, piRNAs) and 60–70 bp (tRNAs, pre-miRNAs, snRNA fragments). Highlighted regions (20-30 bp, 60-70 bp) represent common lengths of EV-enriched RNAs, indicating successful enrichment and trimming accuracy. (**h**) Proportion of Read1 sequences containing UMI-derived technical read-through sequences across multiple tested samples. This panel highlights the significant presence of these artifacts, underscoring the importance of rigorous trimming procedures in EV RNA-seq analyses. (**i**) Mapping statistics for clean reads aligned with STAR, distinguishing uniquely mapped and multi-mapped reads. The substantial fraction of multi-mapping reads underscores the necessity of EVscope’s expectation-maximization (EM) algorithm, which leverages alignment scores and local read coverage to iteratively assign multi-mapped reads accurately. This approach ensures comprehensive and precise RNA quantification critical for downstream analysis of EV RNA-seq data.

### gDNA correction for RNA expression quantification

Previous studies have demonstrated that extracellular vesicles (EVs) contain substantial amounts of cell-free genomic DNA (gDNA), potentially confounding RNA expression analyses. To address this issue, we implemented a gDNA correction strategy during read count quantification. For a typical stranded RNA-seq library where Read2 represents the sense strand, reads aligning to the same strand as the annotated gene are considered potential gDNA contamination. Therefore, for genes on the positive strand, counts from the positive strand were subtracted from counts on the negative strand, and vice-versa for genes on the negative strand. This strand-specific subtraction enables us to more accurately quantify RNA expression within EV samples by effectively distinguishing true RNA signals from contaminating cell-free gDNA. While this approach effectively removes gDNA-derived signals, we acknowledge that it may also remove genuine antisense transcripts. Therefore, this correction should be applied with caution in studies where antisense RNA regulation is a primary focus.

### Curated human RNA annotation for EV transcriptomic profiling

To facilitate accurate and comprehensive quantification of extracellular vesicle (EV) RNA-seq data aligned to the human reference genome (HG38), we curated an extensive annotation dataset covering 20 distinct RNA categories from multiple authoritative sources. Specifically, annotations derived from GENCODE Comprehensive version 45 (Frankish, et al., 2023) included 70,711 entries, comprising protein-coding genes (23,214 confirmed and 1,119 TEC (To Be Experimentally Confirmed) protein-coding genes pending experimental validation), 20,827 long non-coding RNAs (lncRNAs), and several categories of small non-coding RNAs. Regulatory RNAs included 1,945 microRNAs (miRNAs). Structural and processing RNAs were categorized into small nucleolar RNAs (snoRNAs; 1,020), small nuclear RNAs (snRNAs; 2,094), small Cajal body-specific RNAs (scaRNAs; 51), vault RNAs (4), and miscellaneous small non-coding RNAs (miscRNAs; 1,523), encompassing one small cytoplasmic RNA (scRNA), ribozymes (9), and small RNAs (sRNAs; 6). Additionally, canonical Y RNAs (840) and pseudogenic Y RNAs (56) were annotated separately. Ribosomal RNAs included mitochondrial rRNAs (Mt_rRNA; 2) and nuclear rRNAs along with pseudogenes (587). Transfer RNAs annotated using tRNAscan-SE from GENCODE v45 comprised mitochondrial tRNAs (Mt_tRNA; 22) and nuclear tRNAs (648). Immune-related annotations encompassed immunoglobulin (IG) genes (657) and T-cell receptor (TR) genes (317). Pseudogenes were comprehensively represented by 16,398 entries across multiple subclasses. Additionally, we incorporated a gold-standard set of 102,020 piwi-interacting RNAs (piRNAs) annotations from piRBase (Wang, et al., 2022)and retrotransposon annotations from RepeatMasker v4.0.7 (Dfam2.0), obtained from the UCSC genome browser, covering LINEs (1,124,228), ERVs (554,321), and SINEs (1,807,714). The resulting carefully curated RNA annotation dataset contains a total of 3,659,642 RNAs, provides a robust resource for precise quantification and downstream analyses of EV-derived RNA-seq data. This combined annotation file is available at GitHub: (https://github.com/TheDongLab/EVscope) and archived on Zenodo (https://zenodo.org/records/15577789).

### Generation of genomic meta-region annotation from GENCODE v45 (hg38)

Genomic annotations for the human genome (hg38) were obtained from GENCODE v45 (Frankish, et al., 2023). Transcript and gene structures were extracted using GenomicFeatures (Lawrence, et al., 2013) and rtracklayer (Lawrence, et al., 2009). We generated distinct genomic intervals for the following gene-related regions: 5’ untranslated regions (5’ UTR), exons, 3’ untranslated regions (3’UTR), introns, promoters (defined as the region from 1500 bp upstream to 500 bp downstream of transcription start sites, TSS), downstream regions (2 kb downstream of transcript end sites), intergenic regions, and ENCODE blacklist regions (Amemiya, et al., 2019). Briefly, regions were first extracted directly from the transcript database using appropriate functions (e.g., fiveUTRsByTranscript, threeUTRsByTranscript, exons, intronsByTranscript, promoters, flank, and gaps for intergenic regions) and exported to BED format after chromosome-level clipping using known chromosome lengths from UCSC hg38. Subsequently, BEDTools (Quinlan and Hall, 2010) was used to merge overlapping intervals within each feature type to obtain non-redundant intervals (bedtools merge). Finally, feature regions were prioritized as follows: 5’UTR > exon > 3’UTR > intron > promoter > downstream > intergenic. Lower-priority features were sequentially subtracted from higher-priority intervals to ensure mutually exclusive genomic regions (bedtools subtract). ENCODE blacklist regions were obtained separately from the ENCODE portal (Amemiya, et al., 2019). Detailed instructions for downloading all HG38 annotation files are provided on GitHub (https://github.com/TheDongLab/EVscope), and these files are also archived on Zenodo (https://zenodo.org/records/15577789).

### Screening for common human-associated bacterial contaminants

To systematically evaluate potential bacterial contamination, particularly from human-associated bacterial species, we constructed a comprehensive reference database composed of genomic sequences from 240 diverse mycoplasma strains and one Escherichia coli strain. These genome sequences were sourced from the NCBI Genome Assembly database. Included mycoplasma species represent a wide range of clinically relevant strains, such as *Mycoplasma mycoides, Mycoplasma capricolum, Mycoplasma suis*, and *Mycoplasma leachii*, among others. Additionally, an *Escherichia coli* reference genome (GCF_000005845.2_ASM584) was also included in the database. All downloaded genome sequences were compiled into standardized FASTA format files, which were subsequently used to screen RNA-seq datasets for possible bacterial contamination. The complete reference dataset has been made publicly available at our GitHub repository (https://github.com/TheDongLab/EVscope). This database enabled the accurate identification and exclusion of RNA-seq reads originating from common bacterial contaminants, thereby significantly enhancing the reliability and quality of extracellular vesicle (EV) RNA-seq analyses.

### Curation of bulk- and single-cell-derived reference datasets for EV RNA sequencing data deconvolution

Users can utilize customized bulk- or single-cell-derived reference datasets to perform the deconvolution of extracellular vesicle RNA sequencing (EV RNA-seq) data tailored to their specific research objectives. To demonstrate the utility and flexibility of this approach, we provide two carefully curated reference datasets suitable for EV RNA-seq data deconvolution for accurate and robust inference of cell type proportions from bulk gene expression by ARIC (Zhang, et al., 2022). ARIC utilizes a novel two-step marker selection strategy, including component-wise condition number-based feature collinearity elimination and adaptive outlier markers removal. This strategy can systematically identify effective marker genes that ensure a robust and precise weighted υ-SVR-based rare proportion prediction.

First, we assembled tissue-specific gene expression profiles derived from the GTEx V10 dataset, generating average transcript per million (TPM) expression matrices for each of 54 detailed tissue types. These tissues include adipose-subcutaneous, adipose-visceral (omentum), adrenal gland, artery-aorta, artery-coronary, artery-tibial, bladder, brain-amygdala, brain-anterior cingulate cortex (BA24), brain-caudate (basal ganglia), brain-cerebellar hemisphere, brain-cerebellum, brain-cortex, brain-frontal cortex (BA9), brain-hippocampus, brain-hypothalamus, brain-nucleus accumbens (basal ganglia), brain-putamen (basal ganglia), brain-spinal cord (cervical C-1), brain-substantia nigra, breast-mammary tissue, cells-cultured fibroblasts, cells-EBV-transformed lymphocytes, cervix-ectocervix, cervix-endocervix, colon-sigmoid, colon-transverse, esophagus-gastroesophageal junction, esophagus-mucosa, esophagus-muscularis, fallopian tube, heart-atrial appendage, heart-left ventricle, kidney-cortex, kidney-medulla, liver, lung, minor salivary gland, muscle-skeletal, nerve-tibial, ovary, pancreas, pituitary, prostate, skin-not sun exposed (suprapubic), skin-sun exposed (lower leg), small intestine-terminal ileum, spleen, stomach, testis, thyroid, uterus, vagina, and whole blood. Additionally, to facilitate analyses requiring broader tissue categorization, we generated a second set of reference profiles encompassing 30 major tissue groups, including adipose tissue, adrenal gland, bladder, blood, blood vessel, brain, breast, cervix uteri, colon, esophagus, fallopian tube, heart, kidney, liver, lung, muscle, nerve, ovary, pancreas, pituitary, prostate, salivary gland, skin, small intestine, spleen, stomach, testis, thyroid, uterus, and vagina.

Second, to support robust linear regression-based deconvolution specifically for brain-derived EV RNA-seq data, we curated a single-cell reference dataset from the Human Brain Cell Atlas v1.0 (Siletti, et al., 2023). This dataset comprises mean counts per million (CPM)-normalized expression profiles representing 31 distinct superclusters. These superclusters include 10 non-neuronal types (oligodendrocytes, committed oligodendrocyte precursors, astrocytes, Bergmann glia, oligodendrocyte precursors, ependymal cells, choroid plexus cells, fibroblasts, vascular cells, and microglia) and 21 neuronal types (upper rhombic lip-derived cells, splatter cells, lower rhombic lip-derived cells, mammillary body neurons, thalamic excitatory neurons, amygdala excitatory neurons, medium spiny neurons, eccentric medium spiny neurons, miscellaneous neurons, cerebellar inhibitory neurons, midbrain-derived inhibitory neurons, CGE interneurons, LAMP5-LHX6 and chandelier cells, MGE interneurons, deep-layer near-projecting neurons, deep-layer corticothalamic and 6b neurons, hippocampal CA1-3 neurons, upper-layer intratelencephalic neurons, deep-layer intratelencephalic neurons, hippocampal dentate gyrus neurons, and hippocampal CA4 neurons).

